# A Proposed Drought Response Equation Added to the Münch-Horwitz Theory of Phloem Transport

**DOI:** 10.1101/2020.05.01.070995

**Authors:** John D. Goeschl, Lifeng Han

**Affiliations:** Biosystems Research Group, Industrial and Systems Engineering Department, and Soils and Crop Sciences Department, Texas A&M University, College Station, Texas, USA; School of Mathematical and Statistical Sciences, Arizona State University, Tempe, Arizona, USA

**Keywords:** Phloem transport, Turgor pressure, Drought stress, Transport Speed, Concentration

## Abstract

Theoretical and experimental evidence for an effect of sieve tube turgor pressure on the mechanisms of phloem unloading near the root tips during moderate levels of drought stress is reviewed. An additional, simplified equation is proposed relating decreased turgor pressure to decreased rate kinetics of membrane bound transporters. The effect of such a mechanism would be to decrease phloem transport Speed, but increase Concentration and Pressure, and thus prevent or delay negative Pressure in the phloem. Experimental evidence shows this mechanism precedes and exceeds a reduction in stomatal conductance.

## Review of theoretical background

The Münch hypothesis of phloem transport has been expressed mathematically in various model forms by several authors (e.g. Goeschl *et aI*. 1976, Thompson and Holbrook 2003, Hölttä *et al*. 2006, Jensen *et al*. 2011, Payvendi *et al*. 2014). Mathematically consistent models must include equations for at least five dependent variables, (1) the rates of solute Unloading [jU_i_] in sink areas, (2) Osmotic Influx and Efflux [jW_i_] of water through the sieve tube membrane, (3) turgor Pressure [P_i_], (4) transport Speed (i.e. velocity along the Sieve tube) [vS_i_] and (5) solute Concentration [C_i_] along the sieve tube axis. An empirical equation [6] for a sixth variable, (6) Viscosity of the phloem sap (ŋ_i_ assuming only Sucrose at 25° C) was included in the model by Goeschl *et al*. (1976; from Swindells *et al*. 1958) to adjust the effective conductance to the phloem sap along the sieve tube [LS_i_]. The steady-state, algebraic form of these equations for each (i^th^) sieve element (or computational section) are as follows (see default values of the Independent Parameters in Table 1):

**Table 1:** Independent Parameters, values and units for Figure 1,A detailed rationale for estimating the values of Vmax_(Tot)_ is explained in Goeschl (2019), based in part on measurements of phloem behavior using the C-l1 ESW traccr system.

| Independent Parameter | Symbol | Value | Units |
| --- | --- | --- | --- |
| Membrane permeability | $\xi$ | $5 \times 10^{-5}$ | $\text{sec}^{-1} \text{ cm}$ |
| Leaf Apoplastic Water Potential | $\Psi_{(\text{leaf})}$ | -0.4 | MPa |
| Root Apoplastic Water Potential | $\Psi_{(\text{root})}$ | -0.6 | MPa |
| Gas Law Constant | R | 8.3145 | $\text{m}^3 \text{ Pa mol}^{-1} \text{ K}^{-1}$ |
| Reflection Coefficient | $\delta$ | 1 | |
| Conductance of Sieve tube * | LS | * | $\text{cm}^3 \text{ sec}^{-1} \text{ MPa}$ |
| Cross section area of S.T. ** | AX | ** | $\text{cm}^2$ |
| Area of sieve tube membrane *** | AM | *** | $\text{cm}^2$ |
| Total Vmax for each sink **** | $V_{max(tot)}$ | **** | $\text{sec mol}^{-1}$ |
| Concentration for 1/2 v<sub>max</sub> ***** | K<sub>m</sub> | ***** | molar |
| * Calculated for each computational section (Goeschl *et al.* 1976) |  |  |  |
| ** Calculated from ST radius = $1.2 \text{ m}^{-6}$ | | | |
| *** Calculated from ST radius and the length of computational sections = 1 cm |  |  |  |
| **** Negative to indicate Unloading from the sieve tubes |  |  |  |
| ***** Converted to $\text{mol cm}^3$ in the model which uses CGS units | | | |

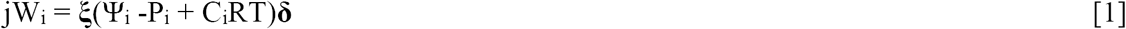

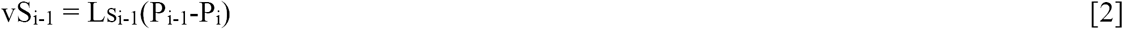

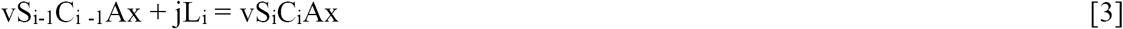

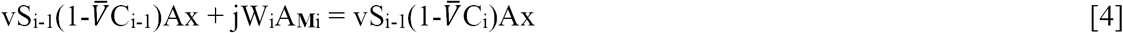

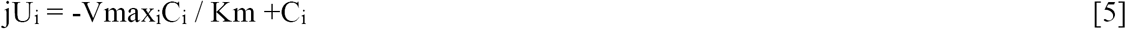

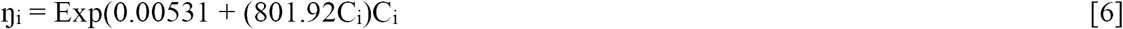

Where **ξ** is membrane permeability, Ψ_i_ is apoplastic water potential, P_i_ is turgor pressure, **δ** is the membrane reflection coefficient, Ls_i_ is the hydraulic conductance of the Sieve Tube, Ax is the cross section area of the Sieve Tube, jL_i_ is the Phloem Loading rate, W is the partial molar volume of sucrose, A_**M**i_ is the surface area of the Sieve Element, and Vmax_i_ and Km are the rate constants of the unloading transporter.

In the course of expressing various predictions of this model (Figure 1), it was observed that uniformly decreasing the value of apoplastic water potential (Ψ_i_) along the axis of a modelled sieve tube (i.e. maintaining the same slope or gradient of water potential) had no effect on the values or patterns of Phloem Unloading, Osmotic Fluxes, Speed, or Concentration, but uniformly lowered the absolute value of Turgor Pressure (Goeschl, 2019).

**Figure 1:**
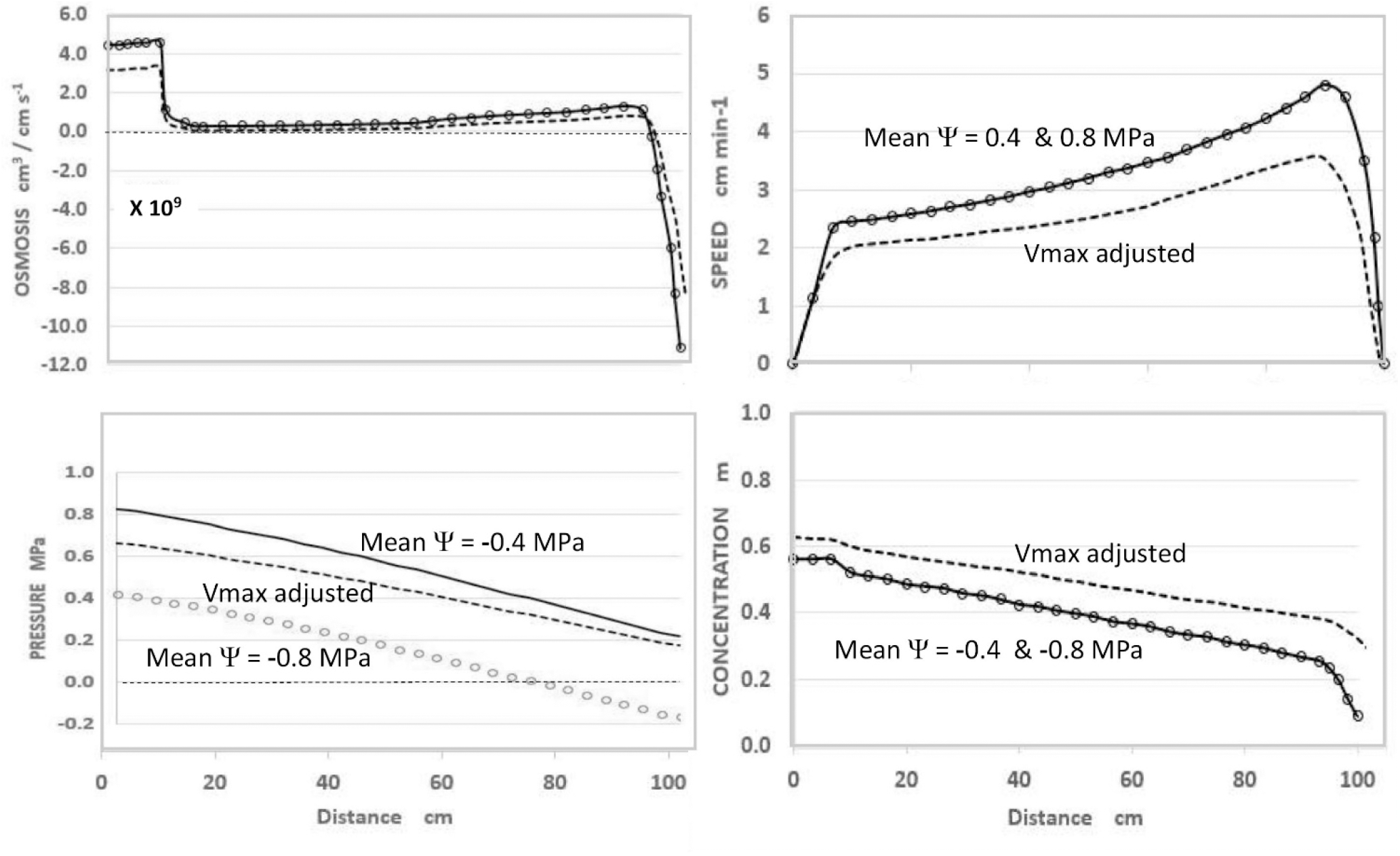
Effects of apoplastic water potential (Ψ) and Effective Unloading Conductance (i.e. EUC assigned as Vmax of the Unloading enzyme system). **Solid Traces** Mean Ψ = -0.4 MPa with Vmax = -1.72 ×10^−10^ mol sec^-1^, **Open Circles** Mean Ψ = -0.8 MPa with Vmax = -1.72 ×10^−10^ mol sec^-1^, **Dashed Traces** Mean Ψ = -0.8 MPa, Vmax adjusted to = -1.35 ×10^−10^ mol sec^-1^ **Note:** Plotted from same data as Goeschl (2019, Figure 7-5) where the term **MPa** should have been **MPa**

As seen in Figure 1, and expected from the Münch hypothesis, the predicted Turgor Pressure always slopes downward from the loading zone to the unloading zone (in this case assumed to be along the root tips). At the chosen values of the independent parameters (Table 1) with a mean water potential of -0.4 MPa the predicted value of Turgor pressure is positive at all points (Figure 1, solid traces). If the mean water potential is lowered to -0.8 MPa, the predicted values of Osmosis, Speed, and Concentration do not change (Figure 1, open circles, superimposed on solid traces). However, Pressure decreases uniformly along the axis (open circles) and reaches negative values approaching the terminal end of the sieve tube. Indeed, one could assign even lower mean values of water potential and the Pressure curve would reach negative values along the entire length of the sieve tube, but the other variables would be unaffected (i.e. would be the same as the solid traces with open circles of Figure 1), so long as the slope of the Pressure curve remained the same.

The question of whether phloem transport in real plants would continue to operate normally with negative pressures was raised by Lang (1974). Among the considerations is that the thin-walled sieve elements in the growing zone of roots are surrounded by newly formed parenchyma and other cells, known to maintain turgor pressures high enough to continue growth in moderate drought stress conditions (Hsaio and Acevedo 1974, Sharp and Davies 1985, Sharp *et al*. 1988, Hsaio and Xu 2000, Hummel *et al*. 1020). It is likely that the sieve elements (possibly including the sieve plate pores) near the root tips would be compressed to smaller diameters, thus restricting the flow of phloem sap. There is also a possibility of plasmolysis under these conditions, as detailed in Goeschl (2019). The question then is whether plants have some mechanism to prevent or minimize the likelihood of negative pressures in the sieve tubes.

One hypothetical mechanism is a reduction in the Effective Unloading Conductance (EUC) of the sieve tubes in the root sinks, e.g. by virtue of a reduction of Vmax (Goeschl, 2019). This could occur in real plants as the sieve tube membranes are compressed, which may alter solute transporter and/or aquaporin protein configurations or expression levels, or membrane electro-potentials, and thus decrease the rate parameters of these mechanisms. This would be consistent with the known effects of turgor pressure on membrane transport kinetics (e.g. Hans *et al*. 1976, Bell and Leigh 1996, Geiger, 2003), on the distribution of photosynthates to various plant organs, especially roots (e.g. Hummel et al. 2010, Lemoine *et al*. 2013) and on related metabolism (Guo *et al*. 2018). Reducing the EUC of the root sinks in a plant with more than one competitive sink, would not necessarily decrease the amount of photosynthates Unloaded into the growing roots. In fact, this could increase (Hummel *et al*. 2010) since reduced EUC would increase solute Concentration throughout the phloem network.

Using the model to predict the effect of reduced EUC in a plant with one hypothetical sink was accomplished by decreasing the collective Vmax_(tot)_ of the hypothetical Unloading Transporters (distributed along the root sink) from 1.72 ×10^−10^ mol sec^-1^ to 1.35 ×10^−10^ mol sec^-1^, while maintaining all other input parameters constant. As seen in Figure 1 (dashed traces), this resulted in a decrease in transport Speed, an increase in Concentration (i.e. reciprocal effects), and substantially increased turgor Pressure to be positive at all points. The relative change in pressure (along with Concentration) was proportionately greater near the root tips. In real plants this would presumably occur gradually and prevent pressure from becoming negative.

## Summary of experimental tests

Experimental measurements were conducted using the Extended Square Wave Carbon-11 Tracer method (^11^C ESW) in individual, live, uninjured, undisturbed Cotton and Corn plants as they were subjected to decreasing water potentials over a period of 4 days (Figure 2, also see Goeschl, 2019, Figures 8-1 and 8-2). The results showed gradual decreases in Transport Speed and increased Concentration, preceding by one or two days, and exceeding the amplitude of decreases in Transpiration (TRANS) and Carbon Exchange Rate (CER, i.e. Photosynthesis) by reduced stomatal conductance.

**Figure 2:**
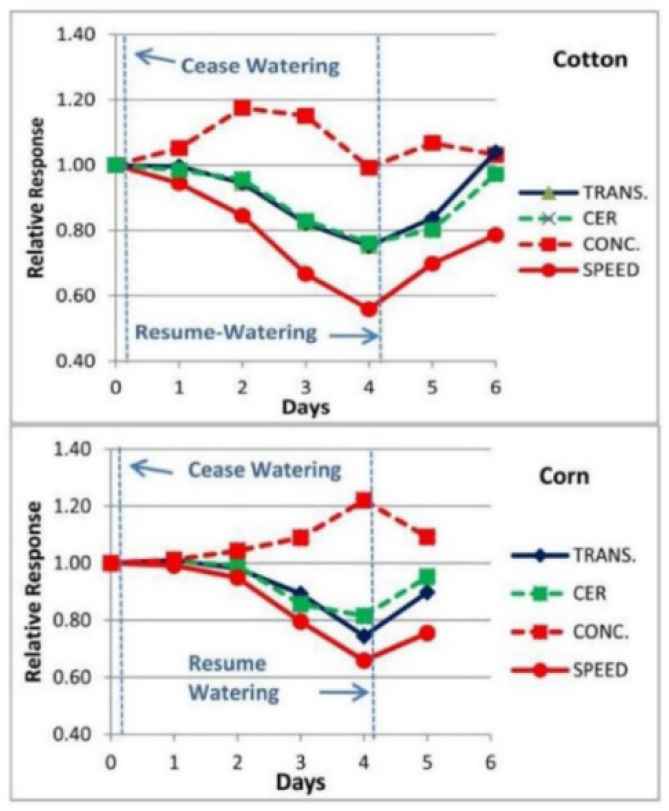
Example of the changes in Phloem Transport Speed and Concentration, (measured experimentally by Carbon-11 Tracer kinetic analysis, Goeschl *et al*. 1988, Goeschl 2019) along with changes in CER and Transpiration of a Cotton and Com plant during a 4 day “drydown” and re-watering experiment of the type conducted by Boyer (1970).

The effects of an immediate restoration of high EUC on transport was also seen on a minute-by-minute basis when moderately drought stressed plants were re-watered (Figure 3 Right). In this case Concentration began to decrease and Speed began to increase immediately, and reached new levels within 40 minutes. This was followed by an increase in Photosynthesis, which led to an increase in both transport Speed and Concentration. The output of a time-dependent version of the Phloem model programmed by Lifeng Han (modified from Smith *et al*. 1980, see Goeschl 2019) closely matched the experimental results.

**Figure 3:**
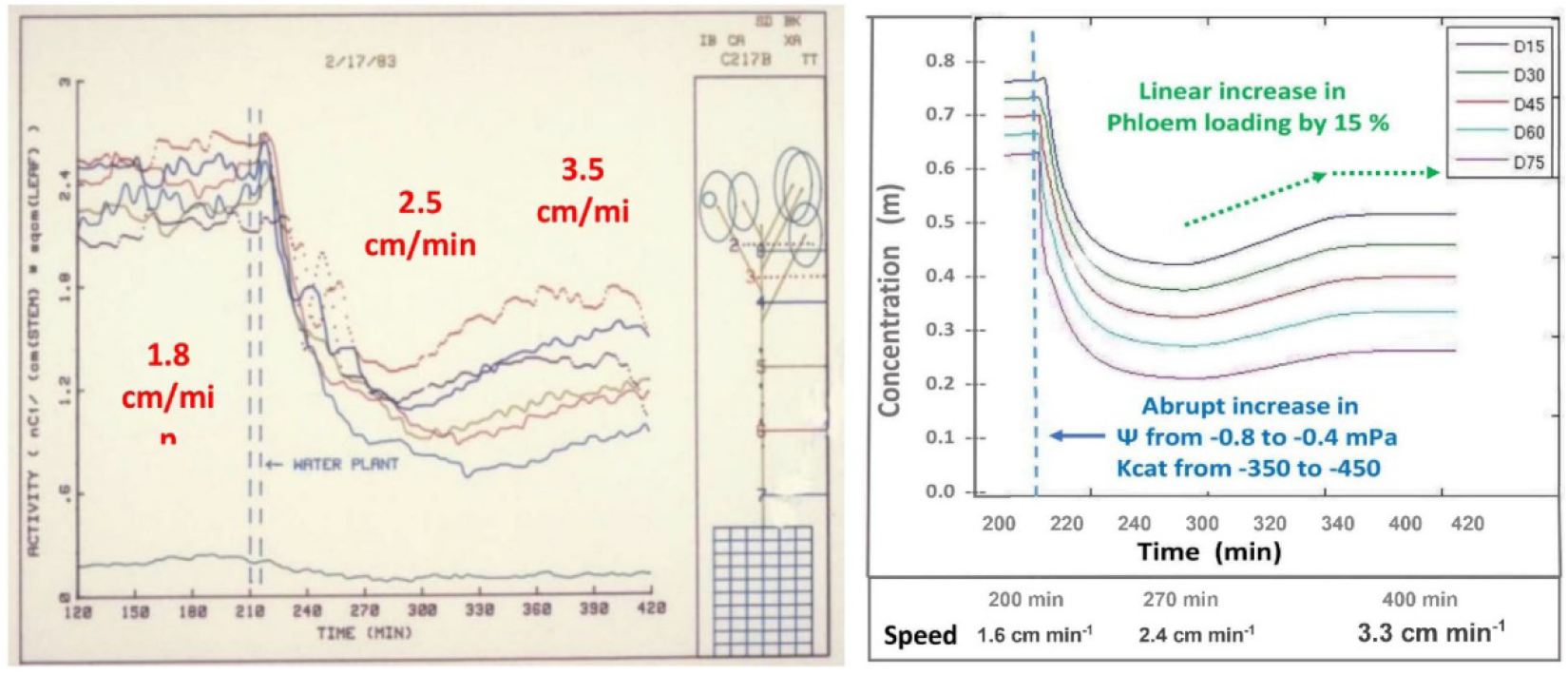
**Left**; Experimental measurement of Carbon-11 activity along the axis of a moderately drought stressed Cotton plant on a minute-by-minute basis before, during, and after rewatering; **Right;** predictions of a time dependent model of the Münch hypothesis subjected to the same hypothetical conditions and unloading response mechanisms proposed to exist the plant.

Mathematical models at the beginning of their development, generally represent the simplest set of assumptions. Such a simple model may adequate to predict the results of the initial assumed circumstances. However, experimental tests may show it to be inadequate to express the effects of additional or changing circumstances. This appears to be the case for the Münch-Horwitz Theory under moderate drought stress conditions (Goeschl, 2019).

If this proposed mechanism is true, and the rate kinetics of the unloading process are altered by the local turgor pressure at appropriate points along the sieve tubes, then the values of one or more of the Independent Parameters of the equation(s) representing the unloading mechanisms (in this hypothetical case the Michaelis-Menten Equation [5]) would become Dependent Variables as a function of Pressure.

Again, starting with the simplest concepts and mathematics one can suggest the following added equation (Equation [7]) where the kinetic parameter Vmax_(max)_ of the membrane bound transporter, or a system of enzyme activities, in the sinks (at the highest likely value of sieve tube turgor Pressure of plants in a growth medium at Field Capacity moisture level) is altered as function of changing Pressures. As a starting point, the following proposed formulation is based on the rationale that the Transporter would be most sensitive as Turgor Pressure approached zero during moderate drought stress levels, and less sensitive as Turgor Pressure approached the levels of relatively unstressed plants. The term Kp is the pressure causing Vmax_(tot)i_ (in the i^th^ computational section) to be ½ of its value at maximal apoplastic water potential and Sieve Tube Pressure. A resulting plot is illustrated in Figure 4.

**Figure 4:**
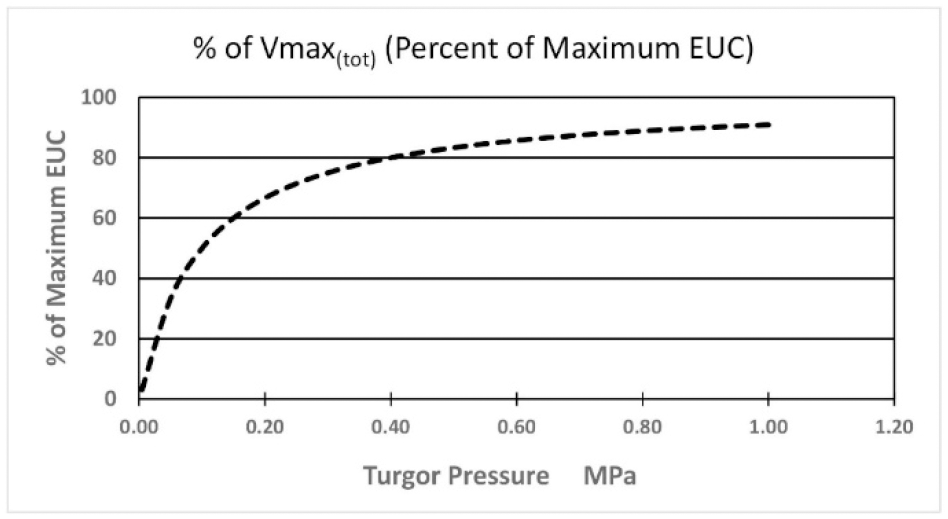
Hypothetical effect of Turgor Presure on the value of Vmax_(tot)_ of the membrane bound Unloading Enzyme System in the root sink.

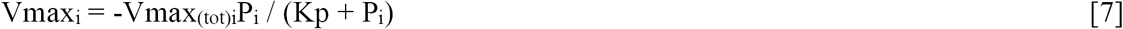

Obtaining realistic values for the parameters of any such equation would not be easy. Again, as a starting point, empirical measurements of values for phloem sap Concentration and Unloading rate in the root zone of real plants, needed to calculate EUC, could be accomplished by combining the ^11^C ESW tracer method (Fares *et al*. 1983, Goeschl *et al*. 1988, Goeschl 2019) with recently developed “Micro-PET” imaging systems (e.g. Cherry *et al*. 1997, Beer *et al*. 2010, Weisenburger *et al*. 2012, Wang *et al*. 2015, and others). This could be performed by quantitative imaging of the final two or three cm length of a growing root, including the root tip as illustrated in Figure 11-4 of Goeschl (2019). Pressure could be measured by probes or estimated on the basis of sieve tube solute Concentration, (calculated from measurements of ^11^C activity during last few minutes of an ^11^C ESW) and the measured values of plant apoplastic water potentials.

Relationships between other Independent Parameters, such as Phloem Loading Rate, Membrane Permeability and Sieve Tube Diameter, in relation to Dependent Variables such as Concentration and Pressure may also exist in real plants and could result in additional equations.

Finally, progress toward a model of carbon flow through the entire plant might be approached by coupling a Phloem Transport model with mechanistic models of Photosynthesis (e.g Zhu *et al*. 2007) and sink metabolism (e.g. Keener *et al*. 1979) to achieve the goals put forward by Demichele *et al*. (1978).

## REFERENCES

Beer, S., M. Struen, T. Hombach, J. Buehler, S. Jahnke, M. Khodaverdi, H. Larur, S. Minwuyelet, C. Parl, and K. Xiemons. (2010) Design and initial performance of PlanTIS: A high resolution positron emission tomograph for plants. Phys. Med. Biol. 55: 635–646

Bell, C.L., and R.A. Leigh. (1996) Differential effects of turgor on sucrose and potassium transport at the tonoplast and plasma membrane of sugar beet storage root tissue. Plant Cell and Environ. 19: 191–200

Cherry, S.R., Y. Shao, R.W. Silverman, K. Medors, S. Siegel, A. Chatziiaonnou, J.W. Young. W.F. Jones, J.C. Moyers, D. Newport, A. Boutefnouchet, T.H. Farquhar, M. Anderson, M.J. Pauluw, D.M. Binkley, R. Nutt, and M.E. Phelps. (1997) MicroPET: A high resolution PET scanner for imaging small animals. IEEE Trans. Nucl. Sci. 44: 1161–1166

Demichele, D.W., P.J.H. Sharpe and J.D. Goeschl. (1978) Toward the engineering of photosynthetic productivity. CRC Critical Reviews of Bioengineering. 3:23–91

Fares, Y., J.D. Goeschl, C.E. Magnuson, C.E. Nelson, B.R. Strain, C.H. Jaeger, and E.G. Bilpuch. (1983) A System for Studying Carbon Allocation in Plants Using 11C-Labeled Carbon Dioxide. Radiocarbon 25:429–439

Geiger, D. (2011) Plant Sucrose Transporters from a Biophysical Point of View. Molecular Plant 4: 395–406.

Goeschl, J.D., C.E. Magnuson, D.W. DeMichele, & P.J.H. Sharpe. (1976) Concentration Dependent Unloading as a Necessary Assumption for a Closed-form Mathematical Model of Osmotically Driven Pressure Flow in Phloem. Plant Physiology 58:556–662,

Goeschl, J.D., Y. Fares, C.E. Magnuson, H.W. Scheld, B.R. Strain, C.H. Jaeger, and C.E. Nelson. (1988) Short-lived isotope kinetics: A window to the inside. In: F.R. Beecher (ed.) Beltsville Symposium in Agricultural Research (11) Research Instrumentation for the 21st Century. Martinus Nijhoff Publishers, (pp. 21–53).

Goeschl, J.D. (2019) Physiological Roles of Phloem Transport: Source-sink Interactions, Drought Stress Responses, Flowering in Plants. Outskirts Press publishers, Denver, CO USA 153 pp.

Guo, R., LX. Shi, Y. Jiao, MX. Li, XL. Zhong, FX. Gu, Q. Liu, X. Xia, and HR. Li. (2018) Metabolic responses to drought stress in the tissues of drought-tolerant and drought-sensitive wheat genotype seedlings. AoB PLANTS 10: ply016; doi: 10.1093/aobpla/ply016

Hans, G., L. Coster, E. Steudle, and U. Zimmermann. (1976) Turgor Pressure Sensing in Plant Cell Membranes. Plant Physiol. 58:636–643.

Hölttä, T., T. Vesala, S. Sevanto, M. Perämäki and E. Nikinmaa. (2006) Modeling xylem and phloem water flows in trees according to cohesion theory and Münch hypothesis. Trees 20:67–78.

Hsiao, T.C. and E. Acevedo. (1974) Plant responses to water deficits, water-use efficiency, and drought resistance. Agric. Meterology 14: 59–84.

Hsiao, T.C. and L.K. Xu. (2000) Sensitivity of growth of roots versus leaves to water stress: Biophysical analysis and relation to water transport. Jour. Expt. Bot. 51: No. 350 WD Special issue pp. 1595–1616.

Hummel, I., F. Pantin, R. Sulpice, M. Piques, G. Rolland, M. Dauzat, A. Christophe, M. Pervent, M. Boutelle, M. Stitt, y. Gibon and B. Muller. (2010) Aribidopsis Plants Acclimate to Water Deficit at Low Cost through Changes of Carbon Usage: An Integrated Perspective Using Growth, Metabolite, Enzyme, and Gene Expression Analysis. Plant Physiol. 154: 357–372

Jensen, K.H., J. Lee, T. Bohr, N.M. Holbrook and M.A. Zwienicki. (2010) Optimality of the Münch mechanism for translocation of sugars in plants. J. R. Soc. Interface doi: 10.1098/rsif.2010.0578

Lang, A. (1974) Tension in the Phloem? Jour. Expt. Bot. 25 (88): 990–994.

Keener, M.E., D.W. Demichele and P.J.H. Sharpe. (1979) Sink Metabolism: A Conceptual Framework for Analysis. Ann. Bot. 44: 659–669.

Lemoine, R., S. La Camera, R. Atanassova, F. Dédaldéchamp, T. Allario, N. Pourtau. J-L. Bonnemain, M. Laloi, P Coutos-Thévenot, L. Maurousset, C. Girousse, P. Lemonnier, J. Parrilla and M. Durand. (2013) Source-to-sink transport of sugar and regulation by environmental factors.

Payvendi, S., K.R. Daly, K.C. Zygalakis and T. Roose. (2014) Mathematical modelling of the phloem: the importance of diffusion on sugar transport at osmotic equilibrium. Bulletin of Mathematical Biology 76: (11) 2834-2865

Sharp, R.E., and W.J. Davies. (1985) Root growth and water uptake by Maize plants in drying soil. Jour. Exptl. Bot. 36: 1441_1456.

Sharp, R.E., W.K. Silk, and T.H. Hsiao. (1988) Growth of the Maize primary root at low water potential: I. Spatial distribution of expansive growth. Plant Physiol. 87: 50–57.

Swindells, J.F., C.F. Snyder, R.C. Hardy and P.E. Golden. (1958) Viscosities of Sucrose Solutions at Various Temperatures: Tables of Recalculated Values. Supplement to National Bureau of Standards Circular 440. July 31, 1958

Thompson, M.V. and N.M. Holbrook. (2003) Application of a Single-solute Non-steady-state Phloem Model to the Study of Long-distance Assimilate Transport. J. Theor. Biol. 220: 419–455

Wang, Q., S. Komarov, A.W. Mathews, K. Li, C. Topp, J.A. O’Sullivan, and Y-C. Tai. (2014) A dedicated high resolution imager for plant sciences. Physics in Medicine and Biology 59: 59–69

Wisenburger, A.G., B. Kross, S. Lee, J. McKisson J.E. McKisson, W. Xi, C. Zorn, C.D. Reid, C.R. Howell, A.S. Crowell, L. Cumberbach, B. Fallin, S. Stolin, and M.F. Smith. (2012) PhytoBeta imager: At positron imager for plant biology. Phys. Med. Biol. 57: 4195–4210.

Zhu, X.G., E de Sturler, and S.P. Long. (2007) Optimizing the Distribution of Resources Between Enzymes of Carbon Metabolism can Dramatically Increase Photosynthetic Rate: A Numerical Simulation Using an Evolutionary Algorithm. Plant Physiology 145: 513–526

